# Demarcating the anatomical structure of the claustrum: piecing together a new functional puddle

**DOI:** 10.1101/179879

**Authors:** Christopher M Dillingham, Bethany E Frost, Maciej M Jankowski, Marie AC Lambert, Shane M O’Mara

## Abstract

**Background:** A major obstacle in the understanding of the functional anatomy of the claustrum has been, and continues to be, the contradiction surrounding its anatomical boundary and, in particular, its rostral (i.e. anterior to striatum), extent. In a recent review we highlighted gene expression-based evidence from the mouse brain which lends weight to the idea that the anatomical boundary of the claustrum does in fact extend rostral to the anterior apex of the striatum. In light of this evidence, in the present study, we have examined the expression of two genes that have previously been identified as having differential expression in the mouse claustrum, with the aim of: 1, establishing the true neuroanatomical boundaries of the rat claustrum; and 2, determining the efficacy of claustral marker expression in the histological verification of claustral electrode placement following electrophysiological recordings in awake behaving rats.

**Methods:** The expression profiles of two genes, crystallin mu (Crym) and guanine nucleotide binding protein (G protein), gamma 2 (Gng2) were assessed immunohistochemically in five male rats (one Wistar Kyoto (WK) and four Lister hooded (LH)). Prior to histological analysis, two of the rats had undergone surgical implantation of tetrodes targeting the putative rostral claustrum.

**Results:** In results that are consistent with those we have previously described in the mouse brain, expression of Crym in the rat brain was highly attenuated, or absent, in the claustrum, demarcating a nuclear boundary that extended considerably beyond the anterior apex of the striatum. In concordance, enriched expression of Gng2 was found in the claustrum, again extending equivalently rostral to the anterior apex of the striatum. The expression of claustral marker genes is a highly effective tool in the verification of electrophysiological electrode placement. Electrophysiological recordings from within this Gng2 and Crym-defined boundary of the rostral claustrum support previous reports of a spatial map in the rostral claustrum.

**Conclusions:** It is now clear that the anatomical boundary of the rat claustrum does in fact extend into more frontal regions of the brain than has previously been asserted. The importance of these findings are considered in the context of the regional specificity of claustral function, anatomical connectivity and electrophysiological properties.

## Introduction

Progress in understanding the complexities of the rodent claustrum has been hindered by a lack of clarity relating to the extent of its anatomical boundary, an issue which is seated in the fact that rodents are lisencephalic and, as such, lack a well defined extreme capsule (a structure that, in gyrencephalic species, provides a clear boundary between the claustrum and the neighbouring cortex; for a recent review, see Dillingham et al. 2017). To overcome the problems that the resulting claustro-cortical continuity presents, a continued focus has been placed on identifying genes that show a differential expression profile in the claustrum relative to surrounding cortical areas. To this end, considerable progress has been made (see Mathur 2014). Crystallin mu (Crym) expression, for instance, is densely expressed in the insular cortex yet is all but absent in the claustrum. Subsequently, Crym expression was fundamental to establishing that that the claustrum is surrounded on all sides by cortex rather than being juxtaposed with the external capsule (Mathur et al. 2009), as was thought previously. In the same study, the nuclear boundary of the claustrum at the level of the striatum was defined using the expression profiles of parvalbumin, cytochrome oxidase and guanine nucleotide binding protein (G protein), gamma 2 (Gng2; Mathur et al., 2009). More recently, Wang et al (2017) compiled a list of 49 genes that were differentially expressed in the mouse claustrum.

Alongside this progression, however, there has been a degree of stagnation in attempts to resolve the issue of whether, or not, the rostral boundary of the claustrum extends beyond the anterior aspect of the striatum. In the seminal work of Mathur et al (2009), the reported absence of parvalbumin and Gng2 expression within the atlas-defined boundary of the rostral claustrum was put forward as evidence for a reassessment of the claustral boundary to one that did not extend beyond the anterior apex of the striatum. Subsequent anatomical and behavioural studies have, for the most part, conformed to the anatomical definition of Mathur et al.

The claustrum is a paired, elongated sheet of grey matter that spans the rostral half of the telencephalon. It has particularly extensive reciprocal cortical connections that show complex topographies in the form of overlapping anterior-posterior gradients of connectivity (Wang et al. 2017). Given the multimodal nature of the claustrum (Remedios et al. 2010) and the likelihood that the separate ‘puddles’ of (presumably functional) connectivity act in concert (Smythies et al. 2014), it is all the more critical that there is consensus in the field relating to its anatomical boundaries.

In a recent review (Dillingham et al., 2017), evidence was put forward that called into question the previous assertion that the claustrum was present only at striatal levels. Using a freely available nucleotide sequence expression mouse brain database (Allen Mouse Brain Atlas; available at: http://mouse.brain-map.org/), the expression of a number of genes that were identified as having differential expression in the claustrum (Mathur et al., 2009; Wang et al., 2017), were assessed. Of 49 genes, the striatal claustrum boundary, delineated either by attenuated expression (e.g. Slit-1, Crym), or enriched expression (e.g. Gng2, Gnb4, latexin), was found to extend considerably rostral to the striatum with its oval cross section situated at the ventrolateral aspect of the forceps minor of the corpus callosum.

In light of this evidence, in the present study, we sought to examine the expression patterns of two genes that have been identified as claustral markers. One of which (crystallin mu; Crym), shows attenuated expression in the claustrum relative to surrounding cortex and the other (Gng2), which shows enriched expression in the claustrum. The expression profiles of these two genes have been reassessed in the rat brain with a particular focus on establishing the boundaries of the rostral extent of the claustrum.

## Methods

### Subjects

A total of five rats were used; four male Lister Hooded rats (Envigo, UK) and one male Wistar Kyoto rat, with pre-procedural weights of between 230-320g.

Of the four LH rats, two were surgically implanted with an electrode bundle containing 28 electrodes arranged in a tetrode formation (platinum-10% iridium; 17 μm thickness; California Fine Wire Ltd., CA, USA), of impedances of between 150-350 kΩ, targeting the claustrum unilaterally. In the same animals, bipolar electrodes (stainless steel, 70 μm thickness) were implanted targeting the CA1 subfield of the septal hippocampus, while electrodes to record myogenic activity were positioned bilaterally beneath neck muscles. All electrodes (32 in total) were fed into a 32-channel microdrive (Axona Ltd., UK).

### Surgical methods

For detailed surgical methods relating to electrode implantation, see Jankowski et al. (2015). Briefly, anaesthesia was induced and maintained with isoflurane (5% and 1–2%, respectively) combined with oxygen (2 L/minute). Animals were then placed in a stereotaxic frame (Kopf, Tujunga, CA, USA) and chloramphenicol eye ointment (Martindale Pharmaceuticals, Romford, UK) was topically applied to the eyes to protect the cornea. Pre-surgical analgesia (Metacam, 1 mg/kg; Boehringer Ingelheim, Germany) and antibiotics (Enrocare; Animal Care Ltd., York, UK) were administered subcutaneously.

The scalp was incised and connective tissue was removed from the skull. With a flat skull, coordinates, derived from the atlas of Paxinos and Watson (2005), for the claustrum (AP +2.5, ML +2.0, DV −5.0) and the CA1 subfield of the septal hippocampus (AP −3.6, ML −3.4, DV −1.9) were marked and craniotomies were made. Tetrodes targeting the claustrum were implanted at an angle 11° laterally in the coronal plane and the hippocampal LFP electrode was implanted at an angle of 18.5° medially in the coronal plane. Both electrodes were anchored to the skull by pre-positioned skull screws and dental cement. In both cases, electromyography wires were positioned bilaterally in between neck muscle compartments before the wound was sutured. Post-operatively, animals were given 10ml glucosaline subcutaneously and then allowed to recover under close observation. In the days following, weight, hydration and activity were monitored twice-daily with a minimum ten day recovery prior to the commencement of electrophysiological recording.

### Electrophysiological recordings

Electrophysiological recordings were performed on two Lister Hooded rats (HippoCla1 and HippoCla2) that had undergone surgical implantation of electrodes (in the rostral claustrum and in CA1 of the septal hippocampus) as part of a separate experiment. The data presented in this study relate only to the properties of spatially tuned units recorded during a behavioural experiment conducted in a bowtie maze (Albasser et al. 2010). Following electrode implantation, rats were habituated to the maze over the course of several days, during which they were trained to push objects, positioned in each arm of the maze, out of the way in order to retrieve a reward (20 mg sucrose pellet; TestDiet, UK). Once both rewards from one half of the maze had been retrieved, a central door was opened and the rats were able to enter the other half of the maze to retrieve the rewards from the other side. Following habituation, during once daily recording sessions, familiar objects were replaced with novel objects (1 per day over the course of 4 days). Following habituation to the novel object set (4 x 15 minute sessions over the course of 4 days), the position of two objects from either side of the maze were switched in order to assess, through time spent exploring objects, whether they were able to detect the change in object position. Prior to behavioural recording, recording sessions were conducted in an open field arena, during which time tetrodes were lowered to their final depth. All units reported were recorded from the final position in order to remove as much uncertainty as possible in terms of histological verification of our recording sites, i.e. there was no requirement for electrode track-reconstruction.

### Immunohistochemistry

Rats were deeply anaesthetised with sodium pentobarbital (Euthanimal) and perfused transcardially with ice cold 0.1M phosphate buffered saline (PBS) followed by 2.5% paraformaldehyde in 0.1M PBS. The brains were removed and postfixed in the same solution for 48 hours before being transferred to a 25% sucrose in 0.1M PBS solution for 1-2 days for cryoprotection. Sections of 40 μm were cut on a cryostat (Leica CM1850) with one 1:4 series mounted directly on to double gelatin subbed microscope slides. Of the remaining 3 x 1:4 series, one was reacted against an anti-Gng2 polyclonal antibody raised in rabbit (Sigma-Aldrich Ireland Ltd; Wicklow, Ireland), while another was reacted against an anti-Crym monoclonal antibody raised in mouse (Novus Biologicals; Abingdon, UK). Briefly, endogenous peroxidases were removed through reaction in a quench solution containing 10% methanol and 0.3% hydrogen peroxide in distilled water. Following washes in PBS and subsequently PBST (0.05% Triton X-100 in 0.1M PBS), the sections were agitated in a 4% solution of normal horse serum in 0.1M PBS for 2 hours. Sections were then transferred to a 1:200 dilution of either anti-Crym or anti-Gng2 in 0.1M PBST with 1% normal horse serum and agitated at 4°C overnight. Following washes in PBST, sections were transferred to a 1:250 dilution of biotinylated horse-anti-mouse IgG (for sections reacted against Crym; Vector Labs, UK) or biotinylated horse-anti-rabbit IgG (for sections reacted against Gng2; Vector Labs, UK) for 2 hours. Sections were then washed in PBST before undergoing signal amplification through incubation in the Vectastain ABC solution (Vector Labs, Peterborough, UK) for 2 hours. Following washes in PBST and subsequently PBS, sections were agitated overnight at 4°C. Immunoreactivity was visualised using the chromagen diamino benzidine (DAB; Vector Labs, Peterborough, UK) and in some cases, signal was intensified with by adding nickel chloride to the DAB solution. Sections were then washed in PBS, mounted, and left to dry at room temperature before being dehydrated in ascending alcohols and coverslipped with DPX mountant (Sigma-Aldrich, Gillingham, UK).

### Microscopy and imagings

An Olympus BX51 upright microscope combined with CellSens acquisition software was used for brightfield microscopy. Images were stitched together using Inkscape (version 2.2.0.0; freely available software available from https://inkscape.org/en/download/), and adjusted in FIJI (‘*fiji is just imageJ’* freely available software available from https://imagej.net/Fiji/Downloads). High magnification photomicrographs in the anatomical figures (1-6) are surface plots in which the z-axis represents relative pixel values (0-255), i.e. areas of low gene expression have lower pixel values than areas of high gene expression.

**Figure 1.**
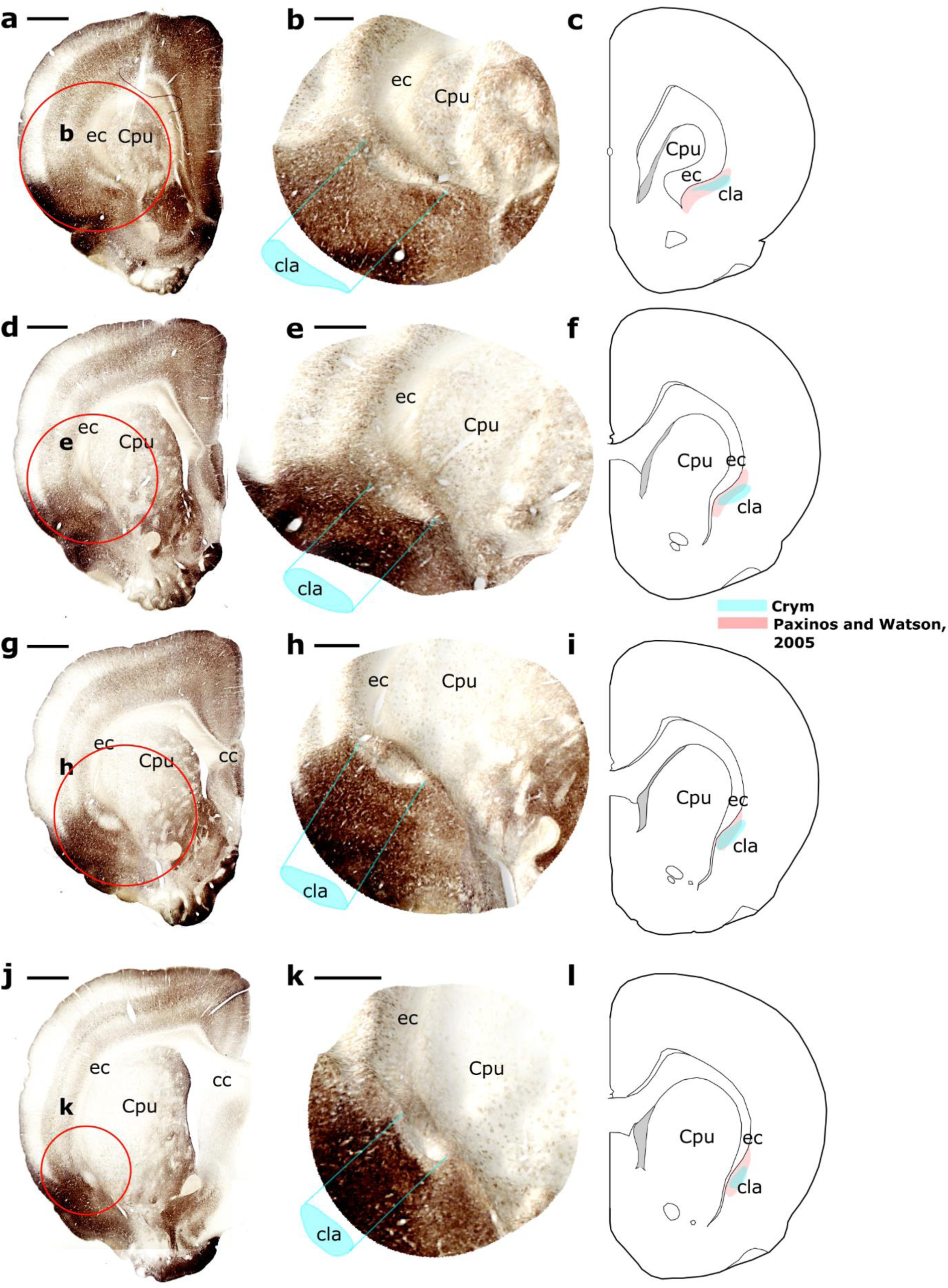
Crystallin mu (Crym) expression is dense in cortical regions, particularly so in the insular cortex, but weak, or absent, in the claustrum. In the striatal claustrum, i.e. at the level of the striatum but rostral to the thalamus, Crym expression clearly delineates the claustral boundary. At more rostral levels (**a-c**), while the medial curavture of the external capsule is greater, the claustrum is more elongated, while further caudally (**d-f**, **g-i** and **j-l**), as the striatum enlarges, the external capsule is pushed laterally and the claustral boundary becomes more vertical and symmetrical in cross-section. At all levels, the medial claustral border is separated from the external capsule by a dense, Crym-immunoreactive, i.e. cortical, region. Red circles in low magnification photomicrographs (e.g. **a**, **d**, **g**, etc.) delineate the region represented by the high magnification surface plots shown in the second column in which contours represent pixel values (e.g. **b**, **e**, **h** and **k**.). Schematic diagrams in the 3^rd^ column (**c**, **f**, **i** and **l**), show the Crym-defined boundary of the claustrum (**turquoise**) superimposed on the atlas-based delineation (**red**; Paxinos and Watson, 2005) at respective anterior-posterior levels. Abbreviations: cc, corpus callosum; cla, claustrum; Cpu, caudate and putamen of the striatum; ec, external capsule. Scale bars (**a**, **d**, **g**, **j**) = 1000 μm; scale bars (**b**, **e**, **h**, **k**) = 500 μm.

### Ethics

Animal husbandry and experimental procedures were carried out in accordance with the European Community directive, 86/609/EC, and the Cruelty to Animals Act, 1876, and was approved by the Comparative Medicine/Bioresources Ethics Committee, Trinity College, Dublin, Ireland, and followed LAST Ireland and international guidelines of good practice.

## Results

### Striatal claustrum

In findings that are consistent with those reported in the mouse by Wang et al. (2017), and in the rat by Mathur et al. (2009), expression of Crym in the rat striatal claustrum was all but absent, contrasting with the dense, predominantly neuropillar, expression in the surrounding deep layer 6 of the insular cortex (as well as the more superficial layers 1-5), which clearly delineate its coronal oval cross section (Fig. 1). Similarly, and consistent with previous reports in the rat (Mathur et al., 2009), expression of Gng2 was found to be enriched in the striatal claustrum but weak, or absent, in surrounding cortical areas (Fig. 2).

**Figure 2.**
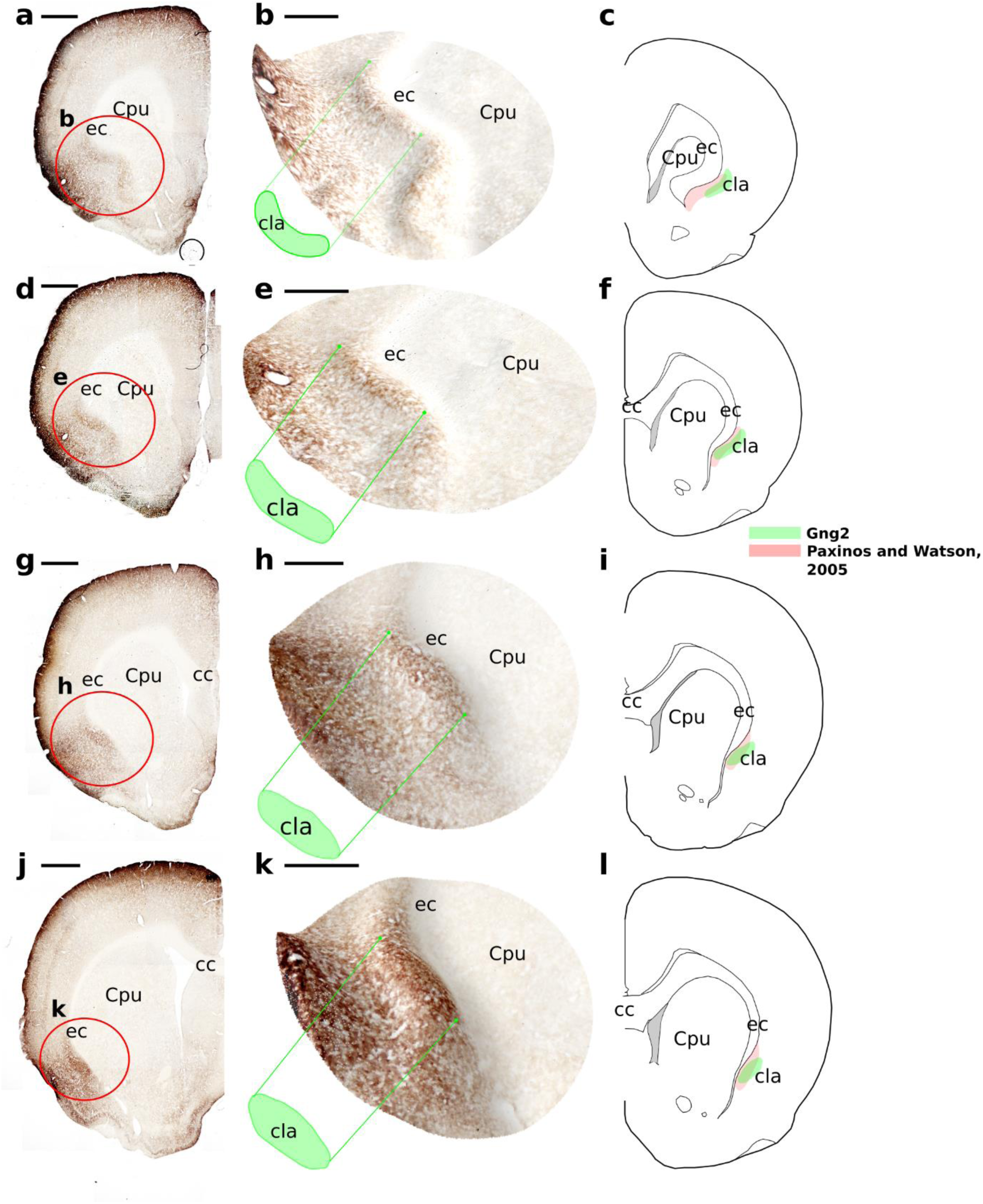
Guanine nucleotide binding protein (G protein), gamma 2 (Gng2) expression is enriched in the claustrum relative to the surrounding cortex. At rostral striatal levels (**a-c**) the claustrum is more elongated, while further caudally (**d-f**, **g-i** and **j-l**), as the striatum enlarges, the claustrum becomes increasingly oval in cross-section. Red circles in low magnification photomicrographs (e.g. **a**, **d**, **g**, etc.) delineate the region represented by the high magnification surface plots shown in the second column in which contours represent pixel values (e.g. **b**, **e**, **h**, etc.). Schematic diagrams in the 3^rd^ column (**c**, **f**, **I**, etc.), show the Gng2-defined boundary of the claustrum (**green**) at each anterior-posterior level, superimposed on the atlas-based delineation (**red**; Paxinos and Watson, 2005). Abbreviations: cc, corpus callosum; cla, claustrum; Cpu, caudate and putamen of the striatum; ec, external capsule. Scale bars (**a**, **d**, **g**, **j**) = 1000 μm; scale bars (**b**, **e**, **h**, **k**) = 500 μm.

### Caudal claustrum

Based upon both Crym and Gng2 expression, the observed caudal extent of the claustrum in the rat was found to be in accordance with the delineation of Paxinos and Watson (2005), where it apexes at the level of the anterior thalamus (Fig. 3). Beyond this anterior-posterior coronal level, regions of Crym attenuation are replaced by continuous cortical enrichment and, similarly, regions of Gng2 enrichment are replaced by continuous cortical attenuation.

**Figure 3.**
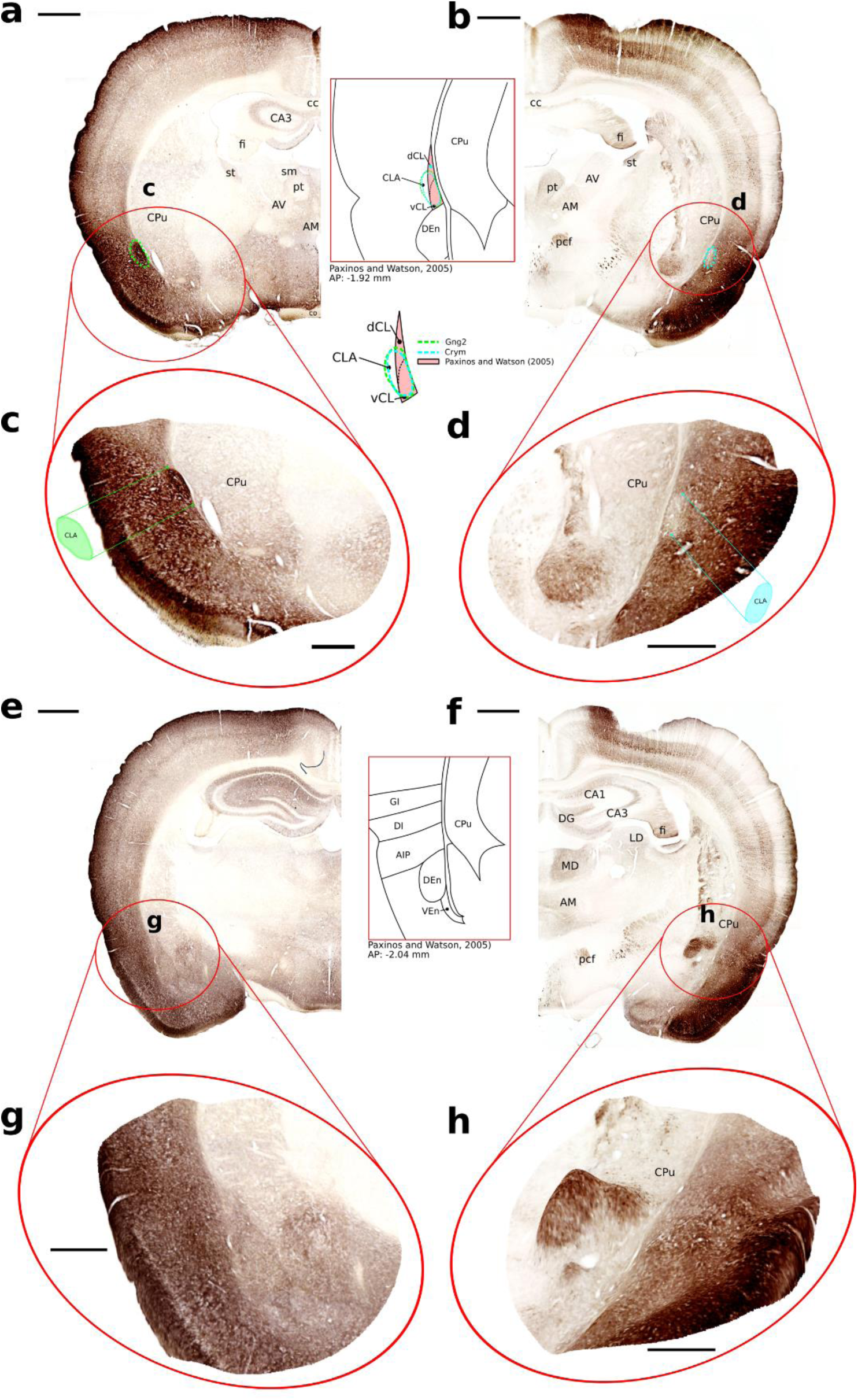
The crystallin mu (Crym) and Guanine nucleotide binding protein (G protein), gamma 2 (Gng2)-defined caudal apex of the claustrum is consistent with the atlas defined delineation, albeit with differences in shape and nomenclature. In the case of both Gng2 (**green**; **a**, **c**) and Crym (**turquoise**; **b**, **d**), the claustrum is still present at the level of the fornix-hippocampal transition. Further caudally, however, at the level of the caudal aspect of the anterior thalamus, neither enrichment in Gng2 expression (**e**, **g**), nor attenutation in Crym expression is observed (**f**, **h**). Red circles in low magnification photomicrographs (**a**-**b** and **e**-**f)** delineate the region represented by the high magnification surface plots in which contours represent pixel values (**c**-**d** and **e**-**f**, respectively). The top inset shows the overlayed claustral boundary as defined by Gng2 (**dashed green**), Crym (**dashed turquoise**) and Paxinos and Watson (**red**; 2005) at the level of the fornix-hippocampal transition. The bottom inset shows a schematic representation of the insular region at a caudal level at which the claustrum is no longer present. Abbreviations: AM, anteromedial thalamic nucleus; AV, anteroventral thalamic nucleus; CA1, CA1 subfield of the hippocampus; CA3, CA3 subfield of the hippocampus; cc, corpus callosum; cla, claustrum; co, optic chiasm; Cpu, caudate and putamen of the striatum; DG, dentate gyrus; ec, external capsule; fi, fimbria of the fornix; LD, laterodorsal thalamic nucleus; MD, mediodorsal thalamic nucleus; pcf, postcommissural fornix; pt, paratenial nucleus; sm, stria medullaris; st, stria terminalis. Scale bars (**a**, **b**, **e**, **f**) = 1000 μm; scale bars (**c**, **d**, **g**, **h**) = 500 μm.

### Rostral claustrum

In concordance with our observations in the mouse brain, the Crym and Gng2 claustral delineation extends rostral to the anterior apex of the striatum in the rat (Figs 4 and 5, respectively). As was observed in the mouse, both Crym and Gng2 expression show the claustrum, at its most rostral extent, to have an oval cross sectional boundary that is situated adjacent to the ventrolateral aspect of the forceps minor of the corpus callosum. As it extends caudally, and as the ventrorbital and lateral orbital cortices subside, the border of the claustrum elongates ventromedially beneath the forceps minor (albeit to a lesser extent to that delineated in the atlases of Paxinos and Watson (1998; 2005; Figs. 4 and 5)). Further caudally, as nucleus accumbens and the striatum enlarge, the external capsule, and with it the claustral body, is pushed laterally, causing the former to reform its oval cross section.

As with the striatal claustrum and its position relative to the external capsule, the rostral claustrum is not juxtaposed with the forceps minor of the corpus callosum. It is apparent from the Crym expression profile that a strip of Crym-immunoreactive neuropil separates the claustrum from the white matter (Fig. 4). Similarly (but less obviously given the continuity of the weakly labelled cortex with the weakly stained forceps minor of the corpus callosum), dense claustral Gng2 expression is separated from the forceps minor by a strip of attenuated Gng2 expression (Fig. 5). The approximate boundary of the rostral claustral border was traced at different anterior-posterior levels based upon Crym and Gng2 expression, independently, and the nuclear boundaries resulting from Crym and Gng2 expression were found to be largely consistent, although the latter showed a boundary that maintained a more regular ovoid shape with a lesser degree of elongation beneath the forceps minor of the corpus callosum (Figs. 4 and 5). While there was, in all instances, a degree of overlap between our gene based delineations and that of Paxinos and Watson (2005), considerable differences were apparent in both the cross-sectional shape as well as in the degree of ventromedial elongation (see Figs. 4 and 5), with our gene-based assessment showing consistently smaller cross-sectional area as well as reduced ventromedial elongation and often extending more dorsolaterally than was expected.

**Figure 4.**
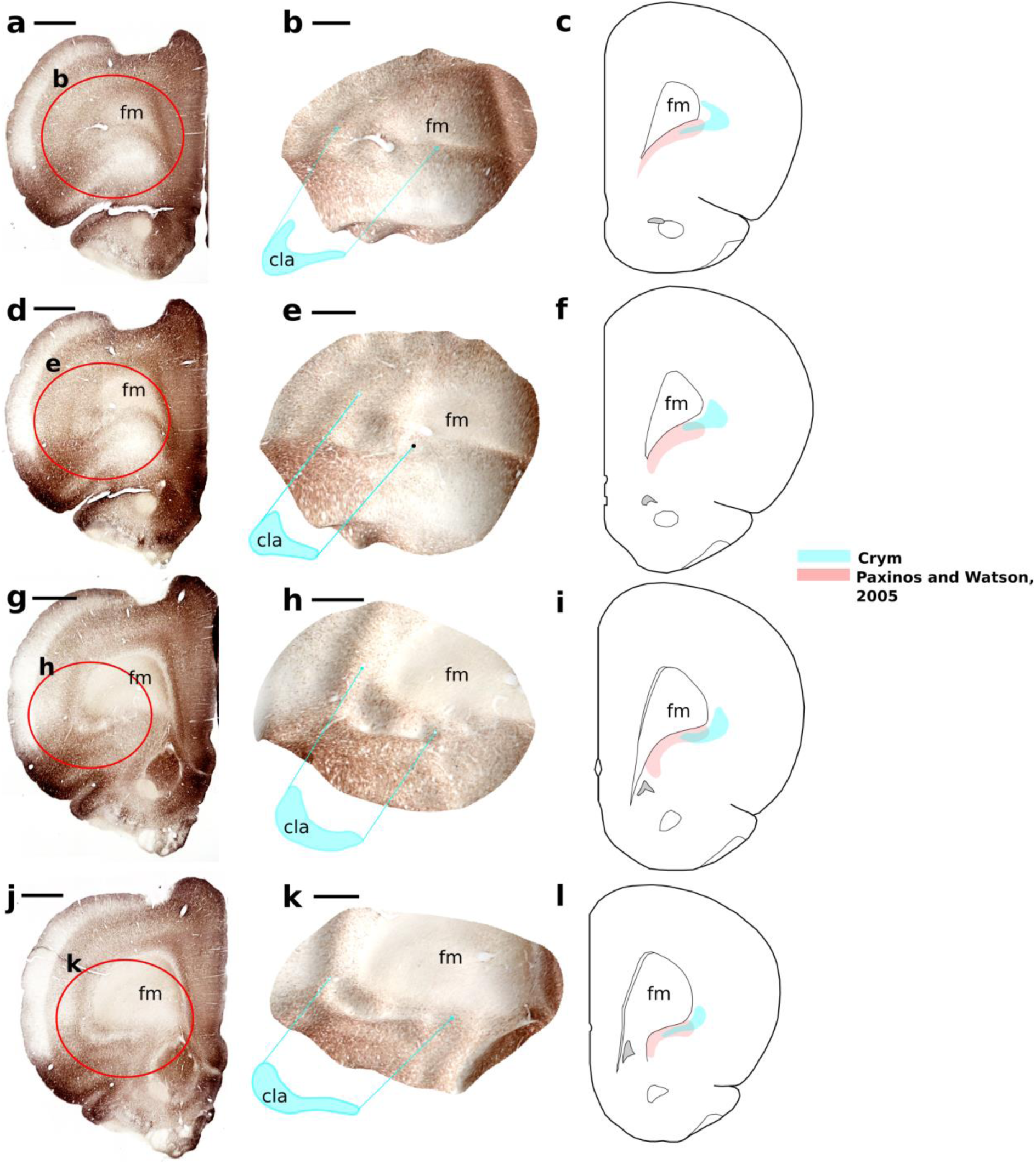
Crystallin mu (Crym) expression delineates the border of the rostral claustrum (i.e. rostral to the anterior apex of the striatum) through an attenuation in immunoreactivity relative to surrounding cortical areas. The rostral most apex of the claustral border, based upon Crym expression, is at the anterior apex of the forceps minor of the corpus callosum (fm). At this rostral level (**a-c**), the coronal cross section of the claustrum is situated at the lateral and ventrolateral border of the forceps minor. It maintains its position here further caudally, extending ventromedially beneath the arch of the forceps minor. The area of attenuated expression that defines the border of the rostral claustrum is continuous with that at striatal levels, as shown in **j**-**k**, which shows a coronal level immediately anterior to the anterior apex of the striatum. Red circles in low magnification photomicrographs (**a**, **d**, **g** and **j**) delineate the region represented by the high magnification surface plots in which contours represent pixel values (**b**, **e**, **h and k**, respectively). Scale bars (**a**, **d**, **g**, **j**) = 1000 μm; scale bars (**b**, **e**, **h**, **k**) = 500 μm.

**Figure 5.**
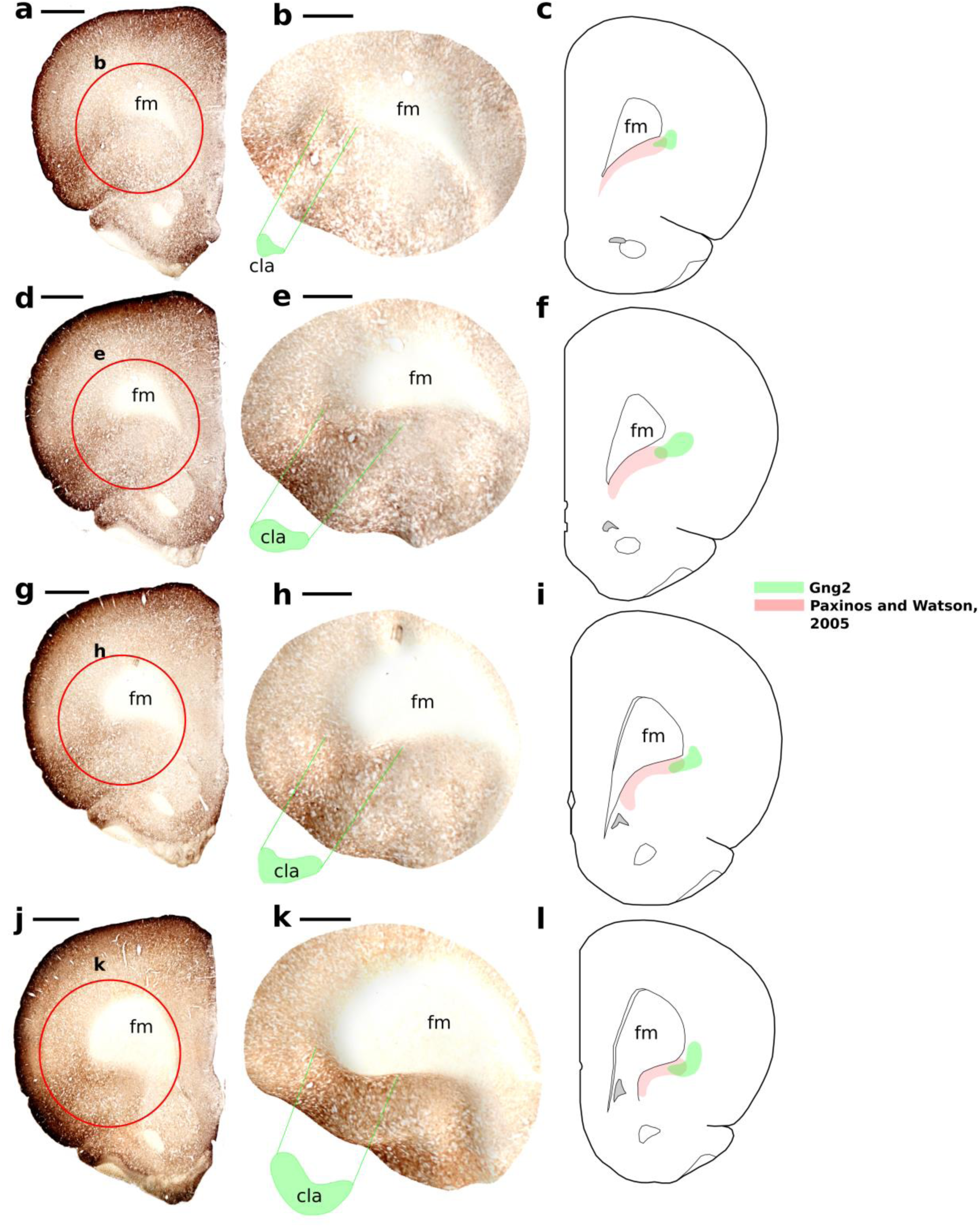
- Guanine nucleotide binding protein (G protein), gamma 2 (Gng2) expression delineates the border of the rostral claustrum (i.e. rostral to the anterior apex of the striatum), through an enrichment in immunoreactivity relative to surrounding cortical areas. Consistent with Crym expression (Fig. 4) the Gng2-defined rostral most apex of the claustral border is at the anterior apex of the forceps minor of the corpus callosum (fm). At this rostral level (**a-c**), the coronal cross section of the claustrum is situated at the lateral and ventrolateral border of the forceps minor. It maintains its position here further caudally, extending ventromedially beneath the arch of the forceps minor. The area of attenuated expression that defines the border of the rostral claustrum is continuous with that at striatal levels, as shown in **j**-**k**, which shows a coronal level immediately anterior to the anterior apex of the striatum. Red circles in low magnification photomicrographs (**a**, **d**, **g** and **j**) delineate the region represented by the high magnification surface plots in which contours represent pixel values (**b**, **e**, **h** and **k**, respectively). Scale bars (**a**, **d**, **g**, **j**) = 1000 μm; scale bars (**b**, **e**, **h**, **k**) = 500 μm.

### Electrode localisation

In two cases, Gng2 and Crym expression was used to validate electrode placement following chronic implantation of tetrodes targeting the claustrum. In the first case (HippoCla1; Fig. 6), Cresyl Violet stained sections showed the electrode track extending ventrally through the longest axis of the forceps minor of the corpus callosum. In more rostral sections in which the electrode track was visible, the electrode recording sites would have been situated in deep orbital cortex, medial to the rostral claustrum, as determined primarily through Crym expression (Fig. 6B). Further caudally, however, as the claustrum elongates ventromedially beneath the arch of the forceps minor, the electrode recording sites were situated within the body of the rostral claustrum as defined by attenuated Crym and enriched Gng2 expression (Fig. 6C). Typically in this case, given that the span of the recording site was both within, and outside the claustrum, i.e. in the deep insular cortex, we would interpret the obtained data with a degree of caution.

**Figure 6.**
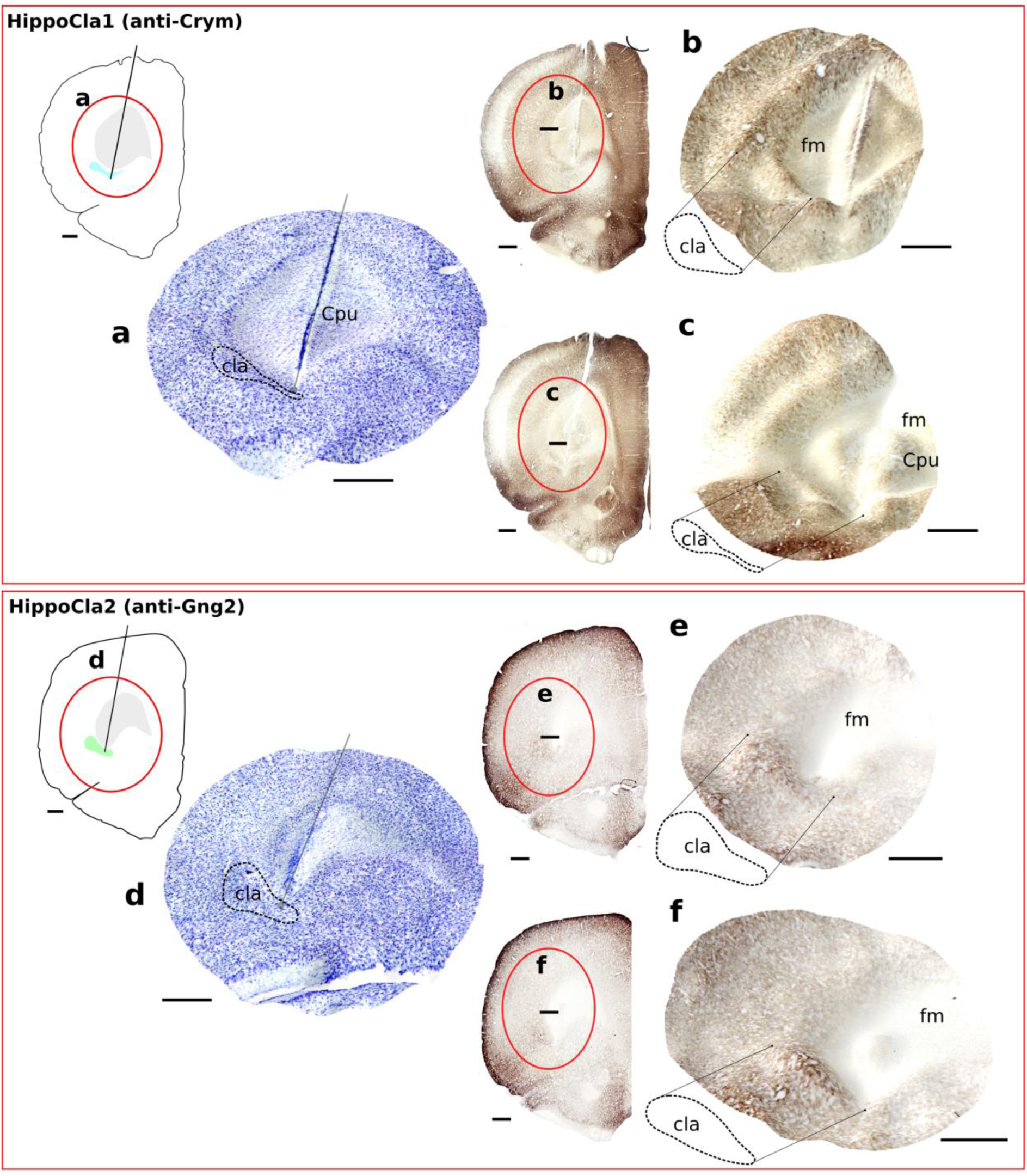
Electrode site verification using standard histological Nissl stain (Cresyl Violet; **A**, **D**), Crystallin mu (Crym) expression and Guanine nucleotide binding protein (G protein), gamma 2 (Gng2) expression, following chronic implantation of tetrodes targeting the rostral claustrum. *Top facet*: The electrode track of HippoCla1 was situated, at rostral levels, medial to the claustrum (**B**) but at more caudal levels, at anterior striatal levels, the electrode track was situated within the claustrum as defined by the region of attenuated Crym expression elongating ventromedially beneath the arch of the forceps minor of the corpus callosum. *Bottom facet:* The electrode track of HippoCla2 was situated at a more rostral levels and in all sections it extended into the region of Gng2 enrichment that defines the boundary of the nucleus. Schematic diagrams in **A** and **D** show the anterior-posterior level of the electrode track, the forceps minor of the corpus callosum (grey) and the claustral cross-section as defined by Crym (**A**, **turquoise**) and Gng2 (**D**, **green**). The red circles in low magnification photomicrographs delineate the region represented by the high magnification surface plots, in which contours represent pixel values respectively). Scale bars = 500 μm.

In the second case (HippoCla2; Fig. 6D-F), the electrode placement was more rostral, with an entry site in a region at the rostral apex of the forceps minor. In all sections, the terminal position of the electrodes were positioned in regions of Gng2 enrichment and Crym attenuation, such that the obtained data could be treated as being derived from the claustrum with a reasonable level of confidence.

### Electrophysiological properties of verified rostral and striatal claustral units

A total of 39 well isolated units were recorded from two rats (6 from HippoCla1 and 33 from HippoCla2), implanted with electrodes in the rostral claustrum (Fig. 7). Given the uncertainty relating to the electrode position of HippoCla1, only data from HippoCla2 will be considered. Of these, 12 units showed spatial tuning, with 7 classified as putative claustral place cells and 5, putative claustral object cells. As is observed with hippocampal place cells in complex environments, e.g. the bow tie maze, some of these putative claustral place cells appeared to exhibit multiple place fields that were consistent and stable across recording days.

**Figure 7.**
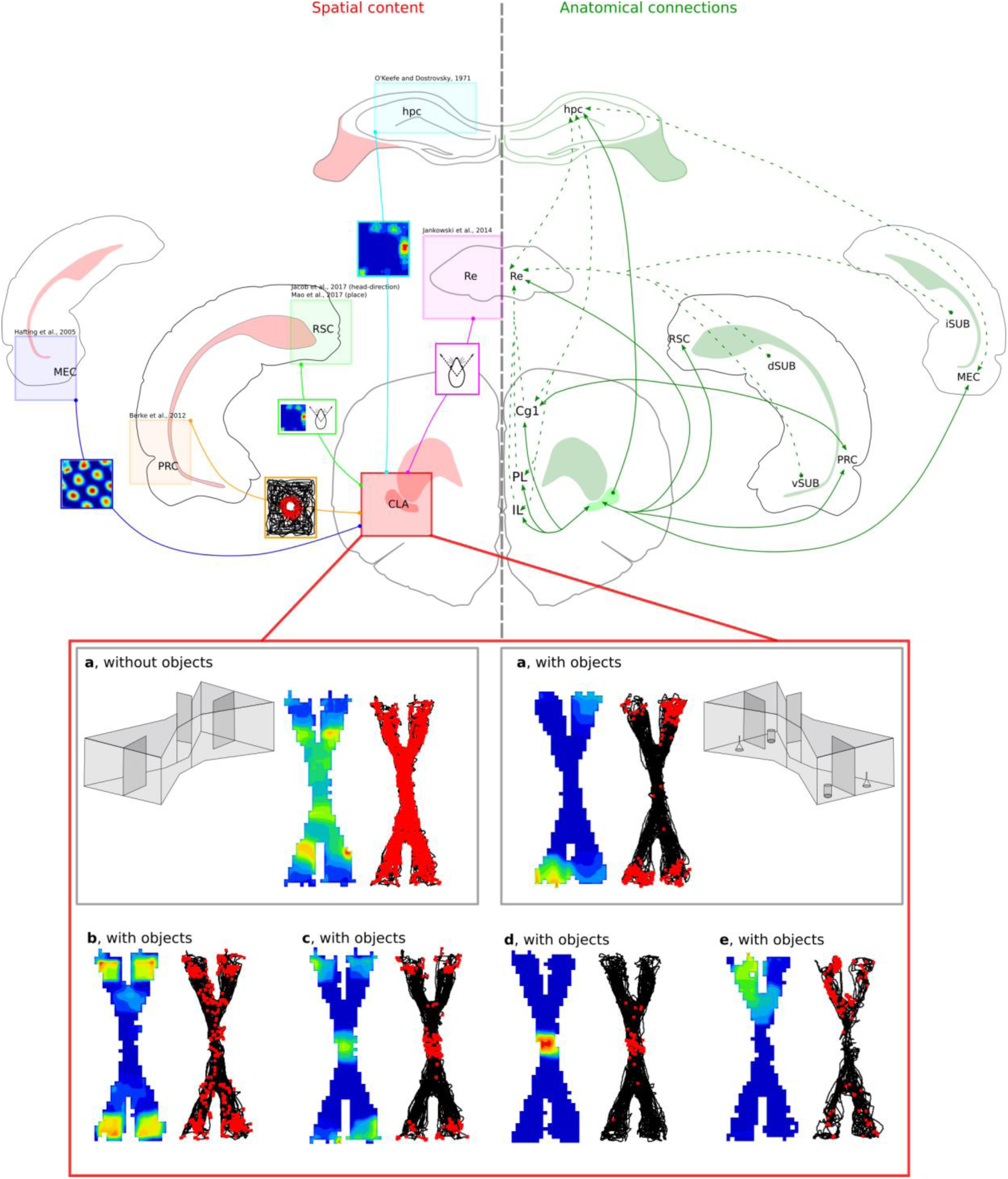
Semi-schematic showing selected anatomical connections (right hand side; **green**) of the rostral claustrum that may underlie, or be related to, its spatial signal. Direct claustral connections (solid green lines) include reciprocal connectivity with medial entorhinal and perirhinal cortices; dense projections to the hippocampus and reciprocal connections with nucleus reuniens. Indirect connectivity that may be implicated in the claustral spatial component revolves around nucleus reuniens, which acts as a critical hub for hippocampal-prefrontal connectivity. Connectivity shown between regions is selected for relevance rather than being exhaustive. Corresponding to these anatomical connections, the corresponding (dominant) spatial content of the regions that are connected to the claustrum are also shown (left hand side; **red**). Medial entorhinal (blue) and perirhinal (orange) cortices are known to house grid and object cells, respectively; both nucleus reuniens and the retrosplenial cortex have a head direction signal, while the hippocampus contains a place signal. In the present study, following stringent electrode localisation using the expression of claustral markers Crystallin mu and Guanine nucleotide binding protein (G protein), gamma 2 (Gng2), several well isolated units were recorded in the course of experiments that showed either place, or object specificity. Unit **A** fired uniformly within the bow tie maze in the absence of objects but when objects were present, unit firing was largely confined to the region in which they were situated, i.e. at the four extremities of the maze. Two further putative claustral object cells are shown, one that appeared to consider the central door an object (**C**), and the other that didn’t (**B**). **D-E** show 2 putative claustral object cells that had stable place field irrespective of the presence, or absence of objects. Abbreviations: CLA, claustrum; Cg1,; dSUB, dorsal subiculum; hpc, hippocampal formation; IL, infralimbic cortex; iSub, intermediate subiculum; MEC, medial entorhinal cortex; PL, prelimbic cortex; PRC, perirhinal cortex; Re, nucleus reuniens; RSC, retrosplenial cortex; vSUB, ventral subiculum. (O’Keefe and Dostrovsky 1971; Hafting et al. 2005; Burke et al. 2012; Jankowski et al. 2014; Jacob et al. 2016; Mao et al. 2017)

Consistent with the reported results of Jankowski and O’Mara (2015), the object cells that were recorded (in HippoCla2), showed distinct activity in the absence of objects with one (Fig. 7; Unit A) showing a consistently high firing rate throughout the maze while another was virtually silent in the absence of objects. In the presence of 4 objects at each extremity of the maze, the previously ubiquitously firing neuron became active only in the areas in which the objects were positioned and virtually silent elsewhere (Fig. 7), while the remaining became active only in the extremities of the maze in which the objects were positioned and remained silent elsewhere (Fig. 7).

## Discussion

In the present study, the expression of two claustral marker genes, Gng2 and Crym, were used to demarcate the boundary of the claustrum of the rat. In results that are largely consistent with those highlighted in the mouse brain (Dillingham et al., 2017), our main finding is that the boundary of the rat claustrum does extend, considerably, beyond the anterior apex of the striatum.

A consensus on the anatomical boundary of the claustrum is key to establishing its functional role and, on a more immediate and practical level, to both the interpretation of, e.g. anatomical evidence, as well as in the verification of electrode placement in electrophysiological studies. Our findings contradict those of Mathur et al (2009) and the reasons for this are not entirely clear. In their study, Mathur et al. also examined the expression of Gng2 and Crym in the rat and as the same antibody and comparable dilutions were used in both studies it is unlikely that the discrepancy is methodological. In their analysis of Gng2 expression, photomicrographs depict an absence of label in the region ventral to the forceps minor of the corpus callosum, i.e. the region defined as the claustrum in the atlas of Paxinos and Watson (2005). It is now apparent that this region is not entirely claustral and, instead, appears to be homogeneous with layer 6 of the insular cortex. Thus, the discrepancy in our findings may be due, in part, to the fact that, at this rostral level, the claustrum is situated more laterally and ventrolaterally to the forceps minor, regions that were not visible in the data presented. The apparent disagreement in our conclusions relating to Crym expression appear to arise from the fact that in the study of Mathur et al., Crym expression was used only to establish that the claustrum is not juxtaposed to the external capsule (as is apparent through its dense cortical expression), but was not used to establish the extent of the overall neuroanatomical boundary of the nucleus. It is likely that if it had been considered in this respect then our findings would have been synonymous. It is worthy of note that while the region of rostral claustrum that we define based upon the expression of Gng2 and Crym shows clear enrichment and attenuation of expression, respectively, the level of attenuation of Crym, in particular, is less accentuated than at striatal levels. On establishing that cortical neurons are positioned between the boundary of the forceps minor of the corpus callosum and the claustrum, Mathur et al (2009) noted that the cortico-claustral boundary is not distinct and that ectopic cortical neurons intermingle with claustral neurons. It would seem, therefore that the extreme rostral extent of the claustrum, i.e. at the level of the anterior apex of the forceps minor, is a claustro-cortical transition zone in which the proportion of ectopic cortical neurons is greater than at more caudal regions.

The unfortunate consequence of these contradictory findings has been the development of a trend in the majority of recent, and in particular anatomical publications, for authors to include a methodological note stating that *analyses of claustral labelling did not extend beyond the most rostral coronal section that contained striatum due to the reported absence of Gng2 expression in these regions*. As a result of this, there is a considerable gap in our anatomical understanding of the claustrum. To our knowledge, only a few studies have directly approached-or included reference to, the rostral claustrum since 2009. Of those that did, the most recent was an excellent anatomical study by Kitanishi and Matsuo (2016) who used anterograde and retrograde pathway tracing to characterise claustro-parahippocampal connectivity. However, rather than relying on atlas-or predefined delineations of the claustrum, they used the expression of claustral markers, e.g. latexin (among others), to define their claustral boundary. Prior to that was an electrophysiological characterisation of the anterior claustrum from our lab (Jankowski et al., 2015), a lesion based behavioural study (Grasby and Talk 2013) as well as a collection of lesion and anatomy based studies looking at the claustrum in the context of epileptogenesis (Zhang et al. 2001; Sheerin et al. 2004).

### Anatomical considerations of the rostral claustrum

In the anatomical study of Zhang et al. (2001), iontophoretic injections of phaseolus vulgaris leucoagglutinin (PHA-L) were placed in the anterior claustrum and while these iontophoretic ejections were guided by the claustral delineation of Paxinos and Watson, (1998; 2005), there is reason to suggest that their anatomical findings are both relevant and important. Take, for instance, that anterograde label resulting from the PHA-L injections in the rostral claustrum resulted in dense fiber and terminal label throughout the striatal body of the claustrum in combination with minimal insular cortical label. This, in the context of the reported findings of Smith and Alloway (2010), which showed that injections of the retrograde tracer Fluorogold into the striatal claustrum resulted in retrograde label throughout the claustrum (through which they demonstrated dense intra-claustral connectivity), suggests that the PHA-L injections of Zhang et al. are likely to have been taken up to a large degree by rostral claustral neurons. Indeed, Zhang et al. also iontophoretically ejected the same tracer as Smith and Alloway (2010; Fluorogold), into the rostral claustrum, and, again, found dense retrograde label in the caudal claustral extent, i.e. showing concordant intra-claustral connectivity. Should this label have been the result of (insular) cortico-claustral, in the case of the PHA-L experiments at least, one would have expected the claustral label to be all but exclusive to the contralateral hemisphere.

Furthermore, the consistency in these reported findings is supported by our findings that, unlike in the mouse, the rostral claustrum does elongate ventromedially beneath the arch of the forceps minor of the corpus callosum, albeit to a lesser degree than that shown in the Paxinos and Watson delineation (Figs. 4 and 5). Additionally, Zhang et al. report that anterograde label in the prefrontal cortex, and indeed all major cortical sites (with some exceptions), was denser in the ipsilateral than the contralateral hemisphere, which is consistent with other reports of claustro-cortical projections being predominanlty ipsilateral. Casting doubt on exclusive claustral uptake, however, it is also reported that retrograde label in the prefrontal cortices was denser in the ipsilateral hemisphere, a finding that is at odds with cortico-claustral projections being predominantly contralateral. Along similar lines, the authors report that retrograde label was present in the contralateral anterior claustrum following unilateral ejection of Fluorogold in the anterior claustrum. While there have been suggestions that inter-claustral connections exist in humans, they have not been reported in the rat.

In light of the above evidence, it is pertinent, with an element of caution, to review the connections reported by Zhang et al., and others, with a view to highlighting those anatomical details relating to the rostral claustrum that are either incomplete or contradictory.

Zhang et al., 2001, did not report anterograde label in the hippocampus following injections of PHA-L into the rostral claustrum. Recently, however, viral tracer injections into dorsal CA1 in the rat showed dense retrograde label in the claustrum (Zhang et al., 2013). In photomicrographs of sagittal sections, dense retrograde label can be seen to extend rostral to the striatum, extending to an anterior-posterior level equivalent to that which we describe here. Thus, it is obvious that clarity is required both in the anatomical nature and indeed the neural functionality of clastro-hippocampal connectivity, particularly so given the putative claustral spatial properties that have been reported (Jankowski and O’Mara, 2015; see below). In terms of parahippocampal connectivity, in a study that utilised, among other approaches, the expression of latexin as a claustral marker, Kitanishi and Matsuo (2016) reported dense reciprocal rostral claustrum-entorhinal cortex connectivity in the mouse. They describe claustral neurons that project to the medial entorhinal cortex (and additionally the prefrontal cortices) as being particularly dense in the anterior claustrum and segregated from sensory and motor cortical projecting neurons in more caudal claustrum. Interestingly, however, no claustro-medial entorhinal cortex projection was reported by Zhang et al (2001) in the rat. Given that claustro-medial entorhinal cortex connections are a likely candidate for involvement in the putative spatial signal in the claustrum, further anatomical studies are required to determine if this is indeed the case in the rat as well or, instead, whether this contradiction is the product of the uncertainty that existed relating to the anatomical boundary of the rostral claustrum.

Although it is unclear which rostral claustral region they are referring to, Smith and Alloway (2010) report that in concordance with the Mathur et al. Gng2 based delineation of the claustrum, the density of retrograde label in the claustrum following tracer depositions in primary motor cortices (M1) declined steeply anterior to the striatum. In line with these findings, Zhang et al. (2001) report that the concentration terminals in M1 was low. Together, these data, and indeed those of Kitanishi and Matsuo (2016; described above) suggest that the gradient of claustro-M1 connectivity may be densest in the striatal claustrum and weaker in the rostral claustrum.

In terms of subcortical connections, Zhang et al. (2001) report rostral claustral projections to include nucleus reuniens, rhomboid nucleus and the mediodorsal thalamic nucleus with retrograde tracer injections suggesting reciprocal connections in most cases. Consistent with the retrograde label in the claustrum following retrograde tracer injection in nucleus reuniens (McKenna and Vertes 2004), Zhang et al. (2001) report anterograde label in nucleus reuniens following PHA-L deposition in rostral claustrum. Similarly, anterograde label in the rostral claustrum following PHA-L injection into nucleus reuniens in a study by Vertes et al. (2006) is supported by the Zhang et al. (2001) report of retrograde label in nucleus reuniens following retrograde tracer deposition (Fluorogold) in the anterior claustrum.

### Electrode localisation with Crym and Gng2 expression

Neuronal population specificity in stimulation studies has advanced greatly within the last decade with the advent of techniques such as optogenetics and DREADDs. Similarly, in terms of electrophysiological recording, the number of regions and neurons that can now be recorded from simultaneously within a single animal is considerable and continues to increase exponentially with time. Electrophysiological characterisation of small and irregularly shaped nuclei such as the claustrum, however, remains very challenging and to do so, with confidence, requires a stringent approach to the verification of electrode placement. In addition to outlining the neuroanatomical extent of the claustrum of the rat, we have shown that utilising differentially expressed claustral markers, e.g. Gng2 and Crym, can provide considerable histological reassurance as well as guard against misleading progress in the path towards understanding the function of the claustrum.

### Electrophysiological considerations of the rostral claustrum

Finally, our evaluation of the anatomical boundaries of the claustrum opened up the possibility that the spatially tuned, putative claustral place and object cells reported by Jankowski and O’Mara (2015) may have been insular/orbital rather than claustral. Instead, with electrodes localised within the Gng2 and Crym defined boundary of the rostral claustrum, twelve of thirty-three recorded units showed a distinct spatial tuning (Fig. 7). Moreover, the very recent independent report of claustro-medial entorhinal cortical interconnectivity being confined to the rostral claustrum in the mouse (Kitanishi and Matsuo 2016; Fig. 7), in amongst more established connectivity, e.g. nucleus reuniens and hippocampus (Fig. 7), would seem to, at least, begin to provide some theoretical basis for our electrophysiological findings. That said, given the difficulties associated with electrode implantation in such an irregularly shaped nucleus, whose borders are known to house ectopic cortical neurons (Mathur et al., 2009), it is not possible to say with certainty that the rostral claustrum exhibits spatial properties. That said, our data do lend weight to this supposition.

## Conclusions

Using the expression profiles of two genes that are widely accepted to be differentially expressed in the striatal claustrum, we report here that, contrary to previous reports, the rostral extent of the claustrum in the rat extends anterior to the rostral apex of the striatum in a manner comparable to that which we described previously in the mouse brain. In addition we have demonstrated the efficacy of such gene markers in the localisation of electrode tracks following electrophysiological recording. Furthermore, electrophysiological data recorded from within the Gng2 and Crym-defined boundary of the rostral claustrum lends support to our original report of spatially sensitive neurons in the claustrum. We have discussed these findings in the context of the known connections of the rostral claustrum, highlighting gaps in our knowledge that have resulted from the lack of clarity on this issue, and proposed anatomical networks that are likely to be related to the observed spatial properties.

